# Changes in the internal genitalia of Hermetia illucens (Diptera: Stratiomyidae) during sexual maturation under a coffee pulp-based diet

**DOI:** 10.64898/2026.08.08.743671

**Authors:** Sergio Rodriguez-Aldana, Dimitri Forero, Pablo Benavides-Machado, Marisol Giraldo-Jaramillo

**Affiliations:** Cenicafé; Universidad Nacional de Colombia, Facultad de Ciencias, Instituto de Ciencias Naturales, sede Bogotá, Bogotá, Colombia

**Keywords:** Black Soldier Fly, coffee pulp, fertilization chamber, gametogenesis, reproductive anatomy

## Abstract

The global expansion of Black Soldier Fly (BSF), *Hermetia illucens* (L.), production systems for organic waste management has increased the need to better understand its reproductive biology in order to optimize mass-rearing programs. In this study, we macroscopically characterized chronological morphological changes in the internal genitalia of adult *H. illucens* reared on a sustainable alternative larval diet based on coffee pulp and corn bran. Morphological assessments and dissections were conducted on female and male reproductive tracts at 3, 6, and 9 days after adult emergence. Overall, the general organization of both reproductive systems was consistent with previous descriptions. A notable observation was the presence of a tripartite fertilization chamber in females, composed of three distinct compartments apparently associated with the three spermathecae. The functional significance of this anatomical organization remains to be determined. Females showed progressively advanced ovarian development and reached the clearest morphological indicators of reproductive maturity at 9 days, while males showed the greatest testicular distention and opacity at the same age. Compared with maturation times reported in previous studies using conventional larval diets, these observations suggest a later pattern of reproductive maturation under the coffee pulp-based diet. The observed differences may be associated with the nutritional composition and carbohydrate-to-protein balance of the larval diet. These results provide chronological and iconographic information that may contribute to the optimization of laboratory rearing protocols and the use of coffee by-products in *H. illucens* bioconversion systems.

## INTRODUCTION

The sustained growth of the global population has driven the intensification of various production systems, generating large volumes of organic waste. Improper management of these residues leads to soil degradation and contamination of water resources (Čičková et al. 2015). Currently, developing sustainable strategies to mitigate this issue is a priority; therefore, the challenge lies in the efficient valorization of different organic residues (Del Hierro et al. 2021).

One of the most documented strategies is the implementation of insects such as the Black Soldier Fly (BSF), *Hermetia illucens* (Linnaeus, 1758), a species native to the Americas with a cosmopolitan distribution (Studt-Solano 2010). This species stands out for its ability to degrade large volumes of organic matter during its larval stage, reaching substrate reduction rates of up to 70% (Diener et al. 2011; Lalander et al. 2015).

Given that the use of *H. illucens* in organic waste transformation is increasingly frequent, it is imperative to deepen its study from various perspectives, particularly its reproductive biology. This knowledge is essential for optimizing large-scale mass-rearing programs aimed at decomposing by-products such as coffee pulp for bioconversion into organic fertilizer. The success of these programs depends on critical factors such as reproductive behavior and larval nutrition. Understanding reproductive morphology and physiology allows for the optimization of management conditions and the identification of periods of higher or lower fecundity, ultimately maximizing progeny production efficiency (Malawey et al. 2019).

According to Lüpold et al. (2020), in some dipterans, especially higher dipterans, the configuration of the reproductive system may show specific adaptations based on sex, age, and evolutionary strategy; factors that directly influence processes such as ovarian maturation, sperm quality, and fertility. Previous studies (Ridley 1988; Lüpold et al. 2020) suggest that adaptations in mating frequency and fecundity allow females of certain hymenopterans and dipterans to mate with multiple males. Likewise, in response to increased sperm competition, males may exhibit larger testes for sperm storage.

In males, longevity plays a decisive role in germ cell production, influencing fertility levels (Papanastasiou et al. 2011). According to Demarco et al. (2014), in dipterans such as *Drosophila melanogaster* Meigen, 1830, spermatogenesis occurs in the testis from the apical end toward the base, allowing the identification of specific zones where stem cells, spermatocytes, spermatogonia, and spermatids are located in different stages of development. A similar configuration was found by Malawey et al. (2019) in *H. illucens* males, where germ cells begin their development in the apical region (germinal zone or “hub”) and mature progressively toward the base in synchronous spermatogenesis within the cysts.

In *H. illucens* females, the ovaries are composed of oocytes within ovarioles that change in shape and coloration with age (Munsch-Masset et al. 2023). These ovaries, of the polytrophic meroistic type, are characterized by ovarioles where each oocyte is associated with nurse cells connected by cytoplasmic bridges. This facilitates synchronous development, allowing maturity to be reached at the time of oviposition (Zhang et al. 2023). Furthermore, the internal morphology allows for the observation of differentiated oviposition chambers, a key aspect in sexual maturity studies.

In holometabolous insects, adult reproductive performance is conditioned by environmental factors such as the availability and quality of nutrients ingested during the larval stage (Lahuatte Vera 2022). In species like *Ceratitis capitata* (Wiedemann, 1824), it has been reported that females fed with protein during the larval phase produce a greater number of mature eggs; conversely, those that did not receive protein tended to mate before completing egg development (Kaspi et al. 2002). In the case of autogenous species like *H. illucens*, females can develop functional oocytes without ingesting protein in their adult phase, utilizing reserves accumulated as larvae (Carpentier et al. 2024), a behavior similar to that observed in *Psorophora (Psorophora) ciliata* (Fabricius, 1794) [as *Culex molestus*] (Kassim et al. 2012). This contrasts with obligatory anautogenous species, such as *Tabanus bromius* Linnaeus, 1758, which require external protein sources (such as blood) to complete oogenesis (Krčmar and Maric 2010).

Another relevant evolutionary strategy in higher dipterans is the development of spermathecae. According to Ward (1993), species like *Scathophaga stercoraria* (Linnaeus, 1758) possess three spermathecae that can ensure cryptic post-copulatory choice, allowing for the storage of a larger amount of sperm from larger males. Given the need to delve into the functioning of the *H. illucens* reproductive system, authors such as Malawey et al. (2019) and Chen et al. (2025) have conducted general descriptions of reproductive anatomy, evaluated spermatogenesis and performed testicular morphometric measurements.

Regarding the female system of *H. illucens*, Tang et al. (2024) and Chen et al. (2025) categorized ovarian development into five stages: previtellogenic development, yolk deposition, egg maturation, peak oviposition, and terminal phase. These studies have provided valuable insights into reproductive dynamics in adults fed diets other than coffee pulp. Considering that larval nutrition can influence reproductive traits in holometabolous insects, this study asked whether a coffee pulp-based diet affects the sexual maturation of *Hermetia illucens*.

We hypothesized that adults reared on coffee pulp would exhibit delayed ovarian and testicular maturation compared to the maturation times previously reported for conventional larval diets. Therefore, the objective of the present study was to macroscopically characterize the morphological changes occurring before, during, and after sexual maturity in the ovaries and testis of adult *H. illucens* at 3, 6, and 9 days of age, whose larvae were fed a coffee pulp-based diet, in order to optimize rearing protocols under laboratory conditions.

## MATERIALS AND METHODS

### Location and Rearing Conditions

The study was conducted at the Entomology Discipline of the National Coffee Research Center (Cenicafé), in Manizales, Caldas, Colombia. *Hermetia illucens* specimens used for the morphological description were reared in a breeding unit under controlled environmental conditions: temperature of 25 °C ± 2, relative humidity of 80% ± 10%, and a 12:12 h (light:dark) photoperiod.

The larval diet consisted of a mixture of coffee pulp, corn bran, sugar, and water, in a 3:1:1 ratio (pulp:bran:sugar) per liter of water. Pupae were placed in conical plastic cages with a 20-liter capacity (dimensions: 80 cm bottom diameter, 90 cm top diameter, and 38 cm total height).

### Collection and conditioning of adults

To ensure the use of newly emerged adults (0 days old), pupae were placed in cages at approximately 8:30 a.m. Subsequently, around 1:00 p.m., the emerged adults were sexed and separated into pairs. Each pair was conditioned in 1-liter conical plastic cages with a fine mesh (tulle) lid (dimensions: 25 cm bottom diameter, 33 cm top diameter, and 15 cm height). Each cage contained 15 pairs of 0-day-old adults, allowing for monitoring and the collection of specimens at specific ages (0, 3, 6, and 9 days) for subsequent dissections.

### Sample Preparation and Dissection

Abdominal scalding and dissections were performed following the protocol described by Sasso Porto et al. (2016).

Adults were cooled for 5 minutes at -18 ± 2 °C, after which the abdomen was separated via a linear cut between the thorax and the abdominal region. The abdomen underwent a maceration process in a water bath with 10% potassium hydroxide (KOH) to remove fatty tissue and facilitate the observation of the reproductive organs, particularly ovarioles and testes. Scalding times were adjusted according to the age of the adults: 55 seconds for 3-day-old individuals, 1 minute and 20 seconds for 6-day-old individuals, and 2 minutes for 9-day-old individuals.

Following scalding, the abdomen was placed in a ventral position. Using micro-scissors, a longitudinal cut was made along the lateral regions, delimiting the junction between the dorsal tergites and ventral sternites to separate the cuticle and expose the internal contents. The cleaning and extraction of reproductive structures were performed using dissection needles, water, and saline solution.

### Documentation and Image Analysis

The obtained structures were mounted on slides using a drop of KY lubricating gel. Observations were conducted using a Carl Zeiss stereomicroscope (Stemi 305 line) and a Carl Zeiss microscope (Primo Star line). Photographic records were captured with a 50 MP camera equipped with a Leica Vario-Summicron lens (24 mm, f/1.9), enabling the documentation of macroscopic morphological differences associated with age in each reproductive structure.

## RESULTS

### Morphology of the female reproductive system of Hermetia illucens

The female reproductive system of *H. illucens* (Fig. 1) is composed of a pair of ovaries (ov) located dorsally within the abdominal cavity. Each ovary is distally connected to a lateral oviduct (lo); both ducts converge into a common oviduct (cod), which constitutes the main pathway for the movement of mature oocytes. The spermathecal complex (s) is formed by three spherical reservoirs. Each spermatheca connects to a rigid rod or capsular duct (rr) that facilitates sperm transit. Continuous with this structure is a valve (v), whose function is to regulate seminal flow toward the ringed channels (rc). Each ringed channel joins an ejector duct (ejd) that opens into the common oviduct, the site where the fertilization of mature oocytes occurs. Finally, the fertilized eggs are conducted toward the terminal portion of the reproductive system, which consists of a fertilization chamber (fc) organized into three differentiated compartments (1, 2, and 3).

**Figure 1.**
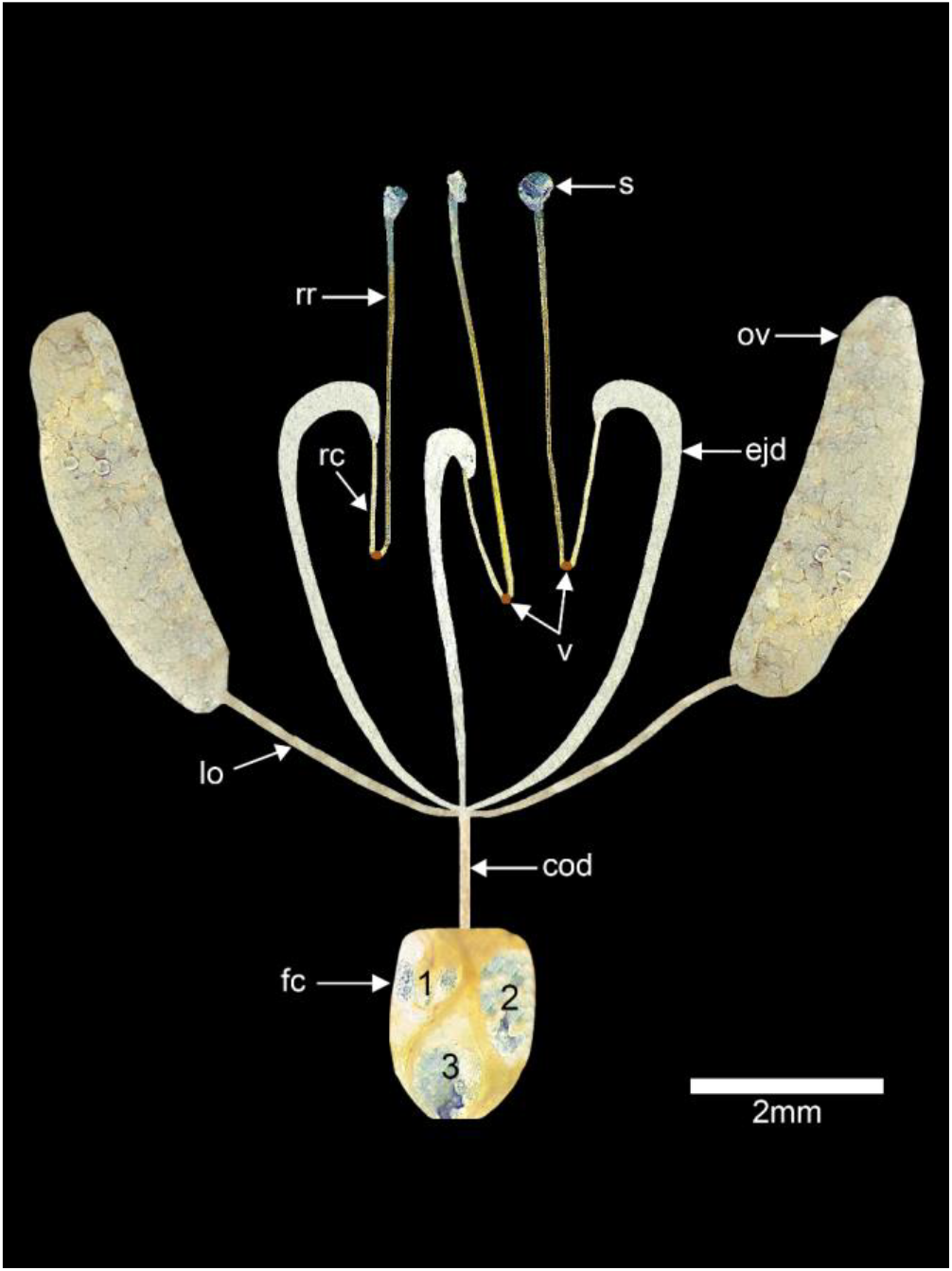
General view of the female reproductive system of *Hermetia illucens*. Indicated are the spermathecae (s), rigid rods (rr), ringed channels (rc), ejector ducts (ejd), valves (v), ovaries (ov), lateral oviducts (lo), common oviduct (cod), and fertilization chamber (fc) with its three compartments (1–2–3).

### Ovarian development in H. illucens females

In 3-day-old females (Fig. 2a), the ovaries are characterized by small, translucent oocytes with an elongated morphology. Initial signs of yolk accumulation are observed, suggesting an active transition from the previtellogenic phase to vitellogenesis. At this stage, the oocytes are densely grouped, reflecting their early state of maturation; overall, the ovaries present a compact appearance and occupy a small portion of the abdominal cavity.

**Fig.2.**
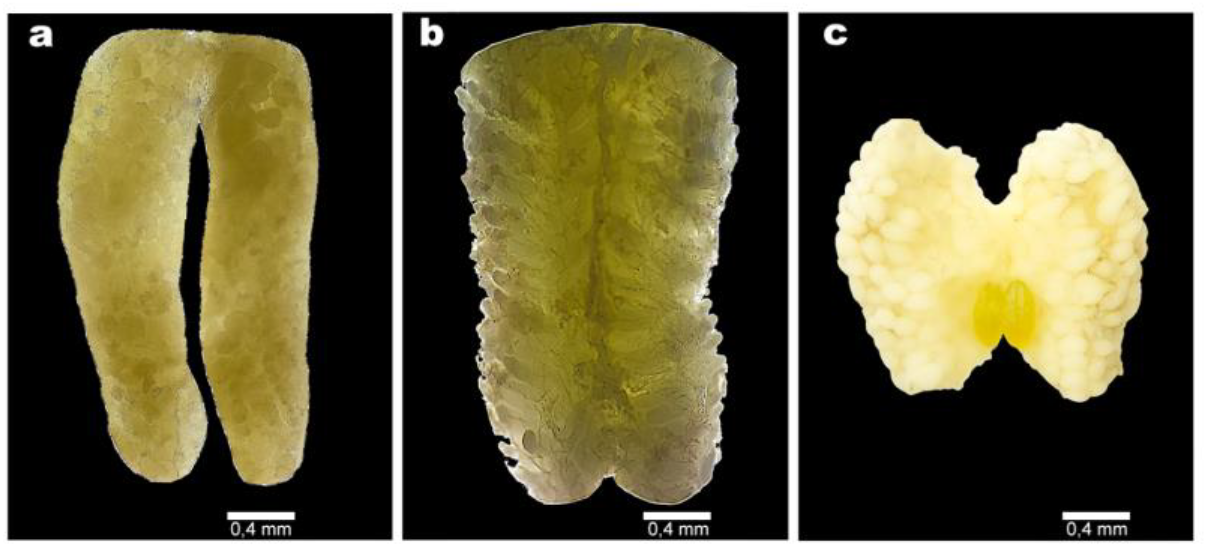
*Hermetia illucens* ovaries at different ages. (a) 3 days: small and translucent oocytes; (b) 6 days: larger and more defined oocytes; (c) 9 days: mature and differentiated oocytes, with yellowish eggs visible in the central region.

By 6 days of age (Fig. 2b), the ovaries of *H. illucens* show more advanced development, with larger and thicker oocytes indicating a progressive vitellogenic phase. A noticeable increase in oocyte volume is evident, and they begin to clearly differentiate from one another, acquiring more defined shapes and showing evident separation within the ovarioles. At this stage, the ovaries are more distended and occupy a significantly larger proportion of the abdomen.

At 9 days of age (Fig. 2c), the ovaries were markedly distended and occupied most of the abdominal cavity. Oocytes were large, individualized, oval, and more opaque than at earlier ages. Yellowish spherical structures corresponding to mature eggs were visible in the central region between the ovaries. These features represented the clearest morphological indicators of reproductive maturity among the ages examined.

### Male reproductive system of H. illucens

The male reproductive system of *H. illucens* (Fig. 3) consists of a pair of testes (t) located dorsally in the abdominal cavity. Each testis gives rise to a vas deferens (vd), which transports sperm toward a common channel (cc) associated with the accessory glands (ag). The common channel is followed by a muscular pump (mp) and a studded tube (sp), which forms part of the ejaculatory duct. The system terminates in the aedeagus (adg), the copulatory organ involved in sperm transfer.

**Figure 3.**
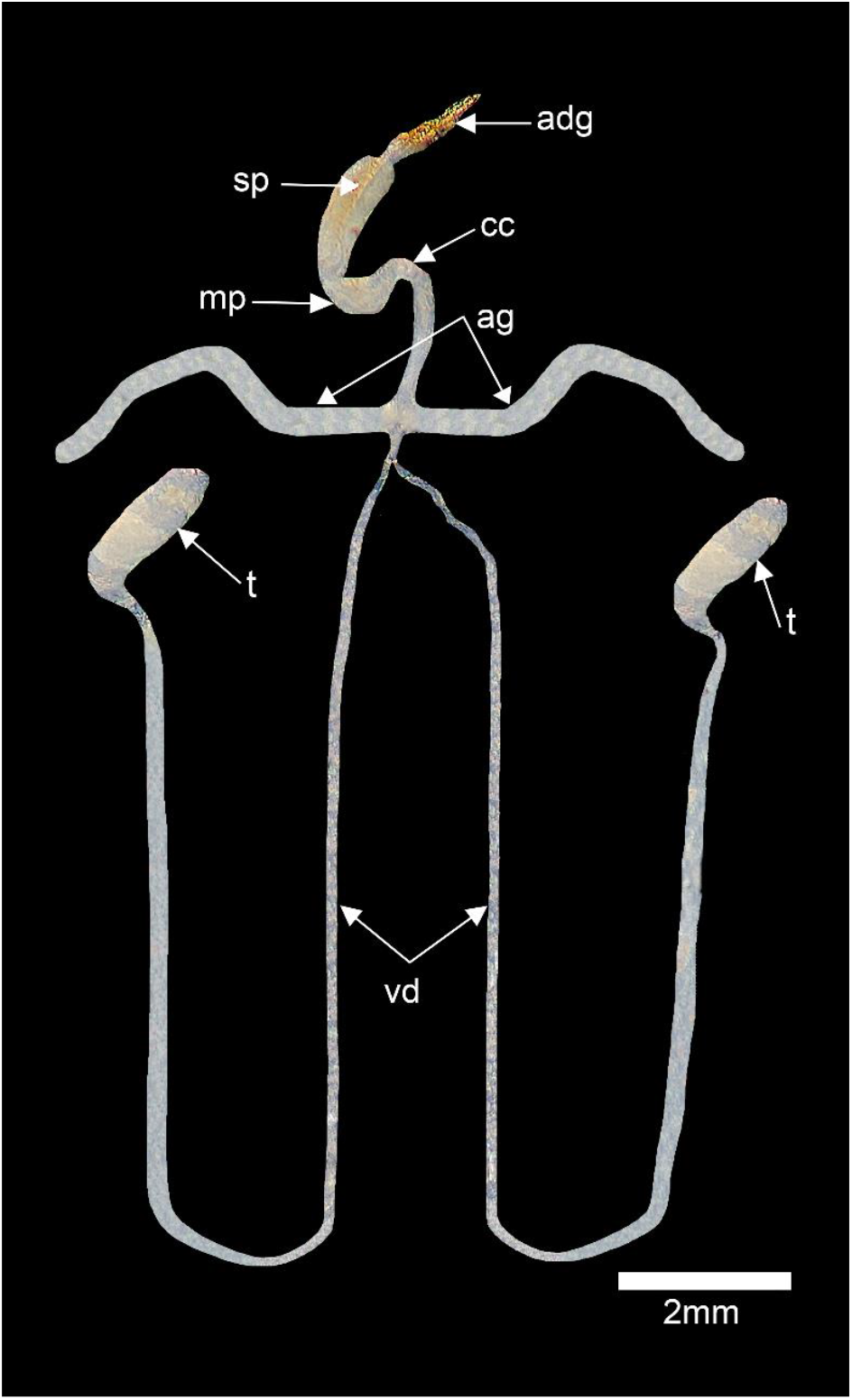
General view of the male reproductive system of *Hermetia illucens*. Indicated are the testes (t), vasa deferentia (vd), accessory glands (ag), common channel (cc), muscular pump (mp), studded tube (sp), and aedeagus (adg).

### Testicular development in H. illucens males

In 3-day-old males (Fig. 4a), the testes were relatively small, translucent, elongated, and slightly distended, consistent with an early stage of reproductive development.

**Figure 4.**
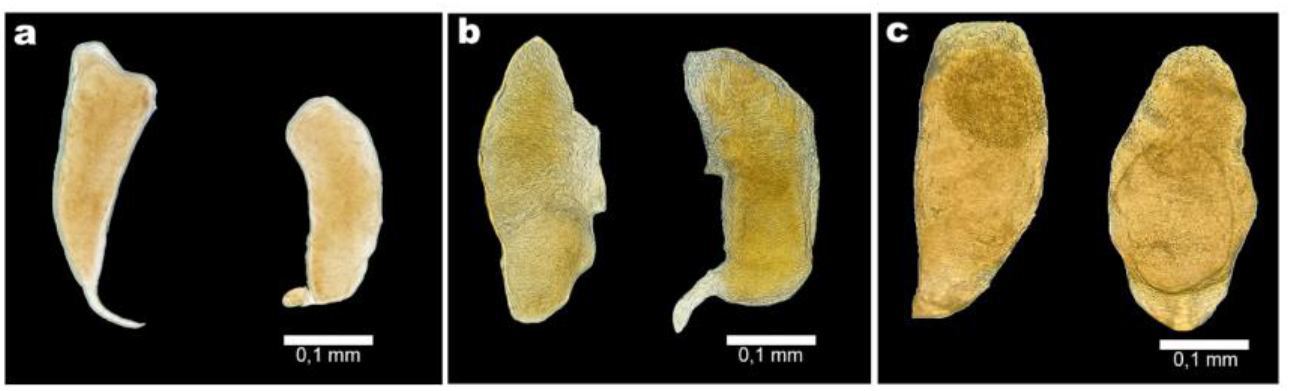
*Hermetia illucens* testes at different ages. (a) 3 days: small, translucent, and slightly distended testes; (b) 6 days: broader testes with a translucent appearance and denser areas; (c) 9 days: distended testes with a more opaque and yellowish appearance.

At 6 days of age (Fig. 4b), the testes were broader and had increased in volume. Although they remained relatively translucent, denser internal regions were visible, indicating continued reproductive development and accumulation of gametes.

At 9 days of age (Fig. 4c), the testes showed their greatest apparent distention and a more opaque, slightly yellowish appearance. These features were consistent with advanced testicular development and greater accumulation of mature gametes.

## DISCUSSION

This study provides a chronological morphological description of the reproductive systems of *H. illucens* reared from larvae on a coffee pulp-based diet. Three observations are particularly relevant: (1) the general organization of the female and male reproductive systems was consistent with previous descriptions; (2) the clearest morphological indicators of reproductive maturity were observed at 9 days of adult age under the conditions evaluated; and (3) a fertilization chamber with three differentiated compartments was observed in females.

The general organization of the *H. illucens* reproductive system agrees with previous reports (Malawey et al. 2019; Munsch-Masset et al. 2023; Tang et al. 2024; Chen et al. 2025). In males, paired testes connect to the vasa deferentia and accessory glands and continue toward the ejaculatory system. In females, the ovaries, lateral and common oviducts, and three spermathecae are consistent with previously described reproductive anatomy.

The three-compartment organization observed in the fertilization chamber is of particular interest. To our knowledge, this specific tripartite organization has not been described previously for *H. illucens*. The three compartments were observed in association with a female reproductive tract containing three spermathecal reservoirs. This anatomical arrangement suggests a possible relationship between sperm-storage structures and the region in which fertilization occurs. However, the present observations are morphological and do not establish direct functional connections between individual spermathecae and compartments, nor do they demonstrate differential sperm storage or utilization.

In other dipterans, the organization of female reproductive structures can be associated with sperm storage and post-copulatory processes. For example, Scolari et al. (2014) described sperm stratification and mixing in *Ceratitis capitata*. Such findings provide a comparative context for considering possible functions of differentiated sperm-storage and fertilization structures in *H. illucens*. Whether the tripartite organization observed here has a role in sperm selection, differential sperm utilization, or other post-copulatory processes remains to be tested experimentally.

The temporal pattern of ovarian and testicular development observed here also warrants consideration. Females showed the clearest morphological indicators of maturity at 9 days, while males exhibited the greatest apparent testicular distention and opacity at the same age. Previous studies have reported earlier reproductive maturation under other larval diets, including maturation or oviposition peaks between approximately 3 and 7 days (Malawey et al. 2019; Tang et al. 2024; Chen et al. 2025). Therefore, the present observations suggest a later pattern of reproductive maturation under the coffee pulp-based diet than that reported in some previous studies.

This comparison should be interpreted cautiously because the present study did not include a conventional-diet control group. Consequently, the data demonstrate the chronological pattern observed under the coffee pulp-based diet but do not by themselves establish that coffee pulp caused the difference in maturation time. Differences among studies may also reflect variation in diet composition, environmental conditions, genetic background, adult management, or other rearing factors. Nevertheless, the nutritional composition of the larval substrate is a biologically plausible factor because protein and carbohydrate balance can influence life-history and reproductive traits in *H. illucens* (Barragán-Fonseca et al. 2019; Zhang and Puniamoorthy 2025).

From a rearing perspective, documenting the age at which reproductive structures show advanced development may be useful for laboratory and mass-rearing systems. The present observations indicate that, under the conditions evaluated, 9-day-old adults showed the clearest morphological signs of reproductive maturation. Future experiments comparing coffee pulp-based and conventional diets under identical environmental conditions, together with direct measures of mating, oviposition, fertility, and sperm storage, would be necessary to determine the extent to which larval nutrition affects reproductive maturation and to establish the function of the tripartite fertilization chamber.

### Conclusions and Implications

Adults of *H. illucens* reared on the coffee pulp-based larval diet exhibited progressive ovarian and testicular development, with the clearest morphological indicators of reproductive maturity observed at 9 days of adult age. This pattern suggests later reproductive maturation than that reported in some previous studies using conventional larval diets, although a direct causal effect of the coffee pulp-based diet cannot be established from the present design. In addition, a tripartite fertilization chamber was observed in females. To our knowledge, this specific organization has not previously been described in *H. illucens*. Its possible relationship with the three spermathecae and with sperm-storage or post-copulatory processes should be investigated experimentally. Together, these observations provide chronological and iconographic information relevant to the reproductive biology of *H. illucens* and may contribute to the optimization of rearing systems based on coffee by-products.

## Notes

### Competing Interest Statement

The authors have declared no competing interest.

